# Physical activity and risks of breast and colorectal cancer: A Mendelian randomization analysis

**DOI:** 10.1101/762484

**Authors:** Nikos Papadimitriou, Niki Dimou, Konstantinos K Tsilidis, Barbara Banbury, Richard M Martin, Sarah J Lewis, Nabila Kazmi, Timothy M Robinson, Demetrius Albanes, Krasimira Aleksandrova, Sonja I Berndt, D Timothy Bishop, Hermann Brenner, Daniel D Buchanan, Bas Bueno-de-Mesquita, Peter T Campbell, Sergi Castellví-Bel, Andrew T Chan, Jenny Chang-Claude, Merete Ellingjord-Dale, Jane C Figueiredo, Steven J Gallinger, Graham G Giles, Edward Giovannucci, Stephen B Gruber, Andrea Gsur, Jochen Hampe, Heather Hampel, Sophia Harlid, Tabitha A Harrison, Michael Hoffmeister, John L Hopper, Li Hsu, José María Huerta, Jeroen R Huyghe, Mark A Jenkins, Temitope O Keku, Tilman Kühn, Carlo La Vecchia, Loic Le Marchand, Christopher I Li, Li Li, Annika Lindblom, Noralane M Lindor, Brigid Lynch, Sanford D Markowitz, Giovanna Masala, Anne M May, Roger Milne, Evelyn Monninkhof, Lorena Moreno, Victor Moreno, Polly A Newcomb, Kenneth Offit, Vittorio Perduca, Paul D P Pharoah, Elizabeth A Platz, John D Potter, Gad Rennert, Elio Riboli, Maria-Jose Sánchez, Stephanie L Schmit, Robert E Schoen, Gianluca Severi, Sabina Sieri, Martha L Slattery, Mingyang Song, Catherine M Tangen, Stephen N Thibodeau, Ruth C Travis, Antonia Trichopoulou, Cornelia M Ulrich, Franzel JB van Duijnhoven, Bethany Van Guelpen, Pavel Vodicka, Emily White, Alicja Wolk, Michael O Woods, Anna H Wu, Ulrike Peters, Marc J Gunter, Neil Murphy

## Abstract

Physical activity has been associated with lower risks of breast and colorectal cancer in epidemiological studies; however, it is unknown if these associations are causal or confounded. In two-sample Mendelian randomization analyses, using summary genetic data from the UK Biobank and GWA consortia, we found that a one standard deviation increment in average acceleration was associated with lower risks of breast cancer (odds ratio [OR]: 0.59, 95% confidence interval [CI]: 0.42 to 0.84, P-value=0.003) and colorectal cancer (OR: 0.66, 95% CI: 0.53 to 0.82, P-value=2*E^-4^). We found similar magnitude inverse associations by breast cancer subtype and by colorectal cancer anatomical site. Our results support a potentially causal relationship between higher physical activity levels and lower risks of breast cancer and colorectal cancer. Based on these data, the promotion of physical activity is probably an effective strategy in the primary prevention of these commonly diagnosed cancers.

**Disclaimer:** Where authors are identified as personnel of the International Agency for Research on Cancer / World Health Organization, the authors alone are responsible for the views expressed in this article and they do not necessarily represent the decisions, policy or views of the International Agency for Research on Cancer / World Health Organization.

## Introduction

Breast and colorectal cancer are two of the most common cancers globally with a combined estimated number of 4 million new cases and 1.5 million deaths in 2018 ^1^. Physical activity is widely promoted along with good nutrition, maintaining a healthy weight, and refraining from smoking, as key components of a healthy lifestyle that contribute to lower risks of several non-communicable diseases such as cardiovascular disease, diabetes, and cancer ^2^.

Epidemiological studies have consistently observed inverse relationships between physical activity and risks of breast and colorectal cancer ^2-5^, but have generally relied on self-report measures of physical activity, which are prone to recall and response biases, and may attenuate ‘true’ associations with disease risk ^6^. More objective methods to measure physical activity, such as accelerometry, have seldom been used in large-scale epidemiological studies, with the UK Biobank being a recent exception, in which ∼100,000 participants wore a wrist accelerometer for 7-days to measure total activity levels ^7^. Epidemiological analyses of these data will provide important new evidence on the link between physical activity and cancer, but these analyses remain vulnerable to other biases of observational epidemiology, such as residual confounding (e.g. low physical activity levels may be correlated with other unfavourable health behaviours) and reverse causality (e.g. preclinical cancer symptoms may have resulted in low physical activity levels).

Mendelian randomization (MR) is an increasingly used tool that uses germline genetic variants as proxies (or instrumental variables) for exposures of interest to enable causal inferences to be made between a potentially modifiable exposure and an outcome ^8^. Unlike traditional observational epidemiology, MR analyses, should be largely free of conventional confounding owing to the random independent assignment of alleles during meiosis ^9^. In addition, there should be no reverse causation, as germline genetic variants are fixed at conception and are consequently unaffected by the disease process ^9^.

We used a two-sample MR framework to examine potential causal associations between objective accelerometer-measured physical activity and risks of breast and colorectal cancer using genetic variants associated with accelerometer-measured physical activity identified from a recent genome-wide association study (GWAS) ^10^. We examined the associations of these genetic variants with risks of breast cancer ^11^ and colorectal cancer ^12^.

## Results

### Mendelian randomization estimates for breast and colorectal cancer

We estimated that a 1 standard deviation (SD) (8.14 milli-gravities) increment in the genetically predicted levels of accelerometer-measured physical activity was associated with a 41% (Odds ratio [OR]: 0.59, 95% confidence interval [CI]: 0.42 to 0.84, P-value=0.003) lower risk of overall breast cancer (Table 2). Similar magnitude inverse associations were found for estrogen receptor positive (ER^+ve^) (OR: 0.53, 95% CI: 0.35 to 0.82, P-value=0.004) and estrogen receptor negative (ER^−ve^) (OR: 0.78, 95% CI: 0.51 to 1.22, P-value=0.27) breast cancer (I^2^=35%; P-heterogeneity by subtype=0.21). There was some evidence of heterogeneity based on Cochran’s Q (P-value<0.05) for the breast cancer analyses; consequently, for these models random effects MR estimates were used (Table 2). MR estimates for each individual single nucleotide polymorphism (SNP) associated with accelerometer-measured physical activity in relation to breast cancer risk are presented in Figure 1.

**Table 1:** Summary information on accelerometer-measured physical activity for the 10 SNPs used as genetic instruments for Mendelian randomization analyses

| SNP | Effect allele | Baseline allele | Chromosome | Position* | Gene | EAF | beta PA<sup>†</sup> | se PA | N<sup>‡</sup> | R | F statistic |
| --- | --- | --- | --- | --- | --- | --- | --- | --- | --- | --- | --- |
| rs12045968 | G | T | 1 | 33225097 | ZNF362 | 0.22 | 0.239 | 0.044 | 91,084 | 0.0003 | 30 |
| rs34517439 | C | A | 1 | 77984833 | DNAJB4 | 0.91 | 0.308 | 0.056 | 91,084 | 0.0003 | 30 |
| rs6775319 | A | T | 3 | 18717009 | LOC105376976 | 0.3 | 0.225 | 0.041 | 91,084 | 0.0003 | 30 |
| rs12522261 | G | A | 5 | 152675265 | LINC01470 | 0.67 | 0.211 | 0.038 | 91,084 | 0.0003 | 31 |
| rs9293503 | T | C | 5 | 88653144 | LINC00461 | 0.88 | 0.329 | 0.059 | 91,084 | 0.0003 | 31 |
| rs11012732 | A | G | 10 | 21541175 | MLLT10 | 0.65 | 0.225 | 0.039 | 91,084 | 0.0004 | 33 |
| rs148193266 | C | A | 11 | 104657953 | RP11-681H10.1 | 0.02 | 0.51 | 0.092 | 91,084 | 0.0003 | 31 |
| rs1550435 | T | C | 15 | 74039044 | PML | 0.53 | 0.2 | 0.037 | 91,084 | 0.0003 | 29 |
| rs55657917 | G | T | 17 | 45767194 | CRHR1 | 0.22 | 0.3 | 0.04 | 91,084 | 0.0006 | 56 |
| rs59499656 | T | A | 18 | 43188344 | RIT2/SYT4 | 0.34 | 0.228 | 0.038 | 91,084 | 0.0004 | 36 |

Abbreviations: EAF: effect allele frequency; PA, physical activity; se: standard error; SNP, single nucleotide polymorphism
\* Position based on GRCh38.p12
<sup>†</sup> The beta coefficients are expressed in milligravities
<sup>‡</sup> N refers to the sample size of the initial GWAS from which the genetic variants were selected

**Table 2.** Mendelian Randomization estimates between accelerometer-measured physical activity and cancer risk.

| Methods | No. Cases | Extended number of SNPs (n=10) |  |  |  | Genome-wide SNPs only (n=2) |  |  |  |
| --- | --- | --- | --- | --- | --- | --- | --- | --- | --- |
|  |  | Estimates (OR)* | 95% CI | P-value | P-value for pleiotropy<sup>†</sup> or heterogeneity<sup>‡</sup> | Estimates (OR)* | 95% CI | P-value | P-value for pleiotropy<sup>†</sup> or heterogeneity<sup>‡</sup> |
| **Breast Cancer** |  |  |  |  |  |  |  |  |  |
| Inverse-variance weighted<sup>§</sup> | 122,977 | 0.59 | 0.42, 0.84 | 0.003 | 6.8×10<sup>-7</sup> | 0.42 | 0.16, 1.16 | 0.09 | 9.4×10<sup>-4</sup> |
| MR-Egger |  | 0.55 | 0.09, 3.20 | 0.50 | 0.9 |  |  |  |  |
| Weighted median |  | 0.76 | 0.59, 0.98 | 0.03 |  |  |  |  |  |
| **ER<sup>+ve</sup> subset** |  |  |  |  |  |  |  |  |  |
| Inverse-variance weighted<sup>§</sup> | 69,501 | 0.53 | 0.35, 0.82 | 0.004 | 0.004 | 0.43 | 0.20, 0.91 | 0.03 | 0.04 |
| MR-Egger |  | 0.61 | 0.07, 5.26 | 0.65 | 0.9 |  |  |  |  |
| Weighted median |  | 0.66 | 0.48, 0.90 | 0.008 |  |  |  |  |  |
| **ER<sup>-ve</sup> subset** |  |  |  |  |  |  |  |  |  |
| Inverse-variance weighted<sup>§</sup> | 21,468 | 0.78 | 0.51, 1.22 | 0.27 | 0.01 | 0.51 | 0.29, 0.89 | 0.02 | 0.11 |
| MR-Egger |  | 0.24 | 0.03, 1.81 | 0.17 | 0.24 |  |  |  |  |
| Weighted median |  | 0.70 | 0.47, 1.04 | 0.08 |  |  |  |  |  |
| **Colorectal Cancer** |  |  |  |  |  |  |  |  |  |
| Inverse-variance weighted | 58,221 | 0.66 | 0.53, 0.82 | 1.9×10<sup>-4</sup> | 0.56 | 0.64 | 0.42, 0.99 | 0.045 | 1 |
| MR-Egger |  | 0.29 | 0.11, 0.80 | 0.016 | 0.10 |  |  |  |  |
| Weighted median |  | 0.67 | 0.50, 0.90 | 0.007 |  |  |  |  |  |
| **Colorectal Cancer in men** |  |  |  |  |  |  |  |  |  |
| Inverse-variance weighted | 30,933 | 0.82 | 0.61, 1.11 | 0.21 | 0.89 | 0.99 | 0.54, 1.81 | 0.98 | 0.31 |
| MR-Egger |  | 0.85 | 0.21, 3.42 | 0.82 | 0.97 |  |  |  |  |
| Weighted median |  | 0.90 | 0.61, 1.33 | 0.60 |  |  |  |  |  |
| **Colorectal Cancer in women** |  |  |  |  |  |  |  |  |  |
| Inverse-variance weighted | 26,848 | 0.54 | 0.40, 0.74 | 1.2×10<sup>-4</sup> | 0.08 | 0.45 | 0.24, 0.84 | 0.012 | 0.26 |
| MR-Egger |  | 0.10 | 0.02, 0.52 | 0.006 | 0.04 |  |  |  |  |
| Weighted median |  | 0.56 | 0.36, 0.88 | 0.01 |  |  |  |  |  |
| **Colon Cancer** |  |  |  |  |  |  |  |  |  |
| Inverse-variance weighted | 30,621 | 0.61 | 0.47, 0.79 | 1.6×10<sup>-4</sup> | 0.57 | 0.57 | 0.34, 0.95 | 0.032 | 0.31 |
| MR-Egger |  | 0.44 | 0.13, 1.45 | 0.18 | 0.59 |  |  |  |  |
| Weighted median |  | 0.55 | 0.39, 0.78 | 0.001 |  |  |  |  |  |
| **Proximal Colon Cancer** |  |  |  |  |  |  |  |  |  |
| Inverse-variance weighted |  | 0.69 | 0.50, 0.97 | 0.03 | 0.83 | 0.75 | 0.39, 1.45 | 0.39 | 0.45 |
| MR-Egger | 13,864 | 0.37 | 0.08, 1.75 | 0.21 | 0.42 |  |  |  |  |
| Weighted median |  | 0.68 | 0.44, 1.04 | 0.08 |  |  |  |  |  |
| **Distal Colon Cancer** |  |  |  |  |  |  |  |  |  |
| Inverse-variance weighted |  | 0.48 | 0.34, 0.67 | 1.9×10<sup>-5</sup> | 0.75 | 0.36 | 0.18, 0.71 | 0.003 | 0.21 |
| MR-Egger | 14,940 | 0.39 | 0.08, 1.89 | 0.24 | 0.8 |  |  |  |  |
| Weighted median |  | 0.5 | 0.31, 0.78 | 0.03 |  |  |  |  |  |
| **Rectal Cancer** |  |  |  |  |  |  |  |  |  |
| Inverse-variance weighted |  | 0.76 | 0.55, 1.07 | 0.12 | 0.64 | 0.91 | 0.47, 1.77 | 0.78 | 0.91 |
| MR-Egger | 15,859 | 0.37 | 0.08, 1.75 | 0.21 | 0.46 |  |  |  |  |
| Weighted median |  | 0.91 | 0.58, 1.42 | 0.69 |  |  |  |  |  |

Abbreviations: CI, confidence intervals; MR: Mendelian Randomization; OR: odds ratio; SNPs: Single nucleotide polymorphisms
\* The estimates correspond to a standard deviation increase in physical activity
† P-value or pleiotropy based on MR-Egger intercept
‡ P-value for heterogeneity based on Q statistic
§ The estimates were derived from a random-effects model due to the presence of heterogeneity based on Cochran's Q statistic

**Figure 1:**
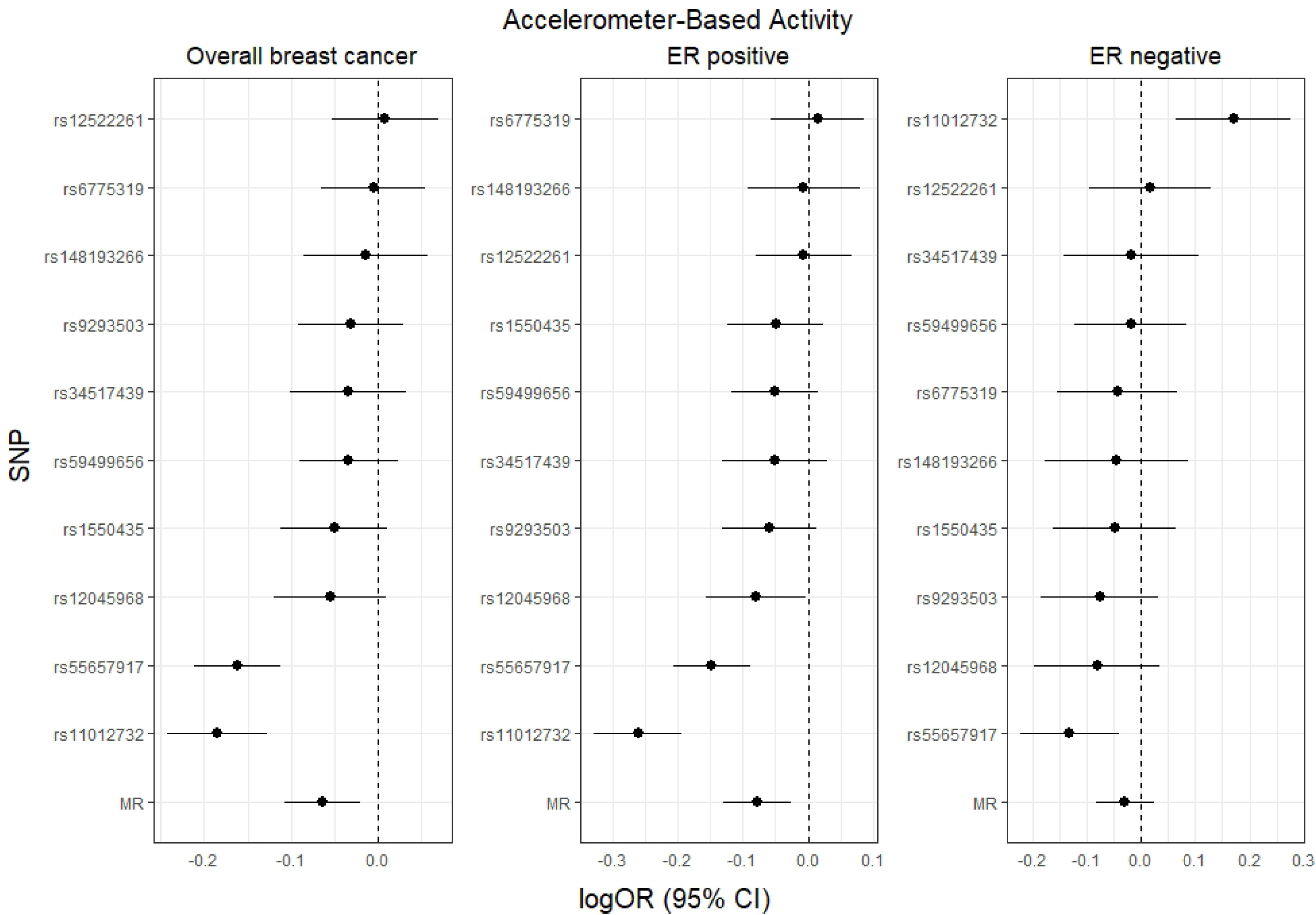
Mendelian randomization analysis for individual SNPs associated with accelerometer-measured physical activity in relation to breast cancer risk. The x axis corresponds to a log OR per one unit increase in the physical activity based on the average acceleration (milli-gravities). The Mendelian randomization (MR) result corresponds to a random effects model due to heterogeneity across the genetic instruments. logOR = log odds ratio. 95% CI = 95% confidence interval. SNP = single nucleotide polymorphism.

For colorectal cancer, a 1 SD increment in accelerometer-measured physical activity level was associated with a 34% (OR: 0.66, 95% CI: 0.53 to 0.82, P-value=1.9*10^−4^) lower risk. The estimated effect size was stronger for women (OR: 0.54, 95% CI: 0.40 to 0.74, P-value=1.2*10^−4^) than men (OR: 0.82, 95% CI: 0.61 to 1.11, P-value=0.21), although this heterogeneity did not meet the threshold of significance (I^2^=72%; P-heterogeneity by sex=0.06). For colorectal subsite analyses, accelerometer-measured physical activity levels were inversely associated with risks of colon cancer (OR per 1 SD increment OR: 0.61, 95% CI: 0.47 to 0.79, P-value=2*10^−4^) and rectal cancer (OR: 0.76, 95% CI: 0.55 to 1.07, P-value=0.12). MR estimates for each individual SNP associated with accelerometer-measured physical activity in relation to breast cancer risk are presented in Figure 2 and Supplementary fig 1.

**Figure 2:**
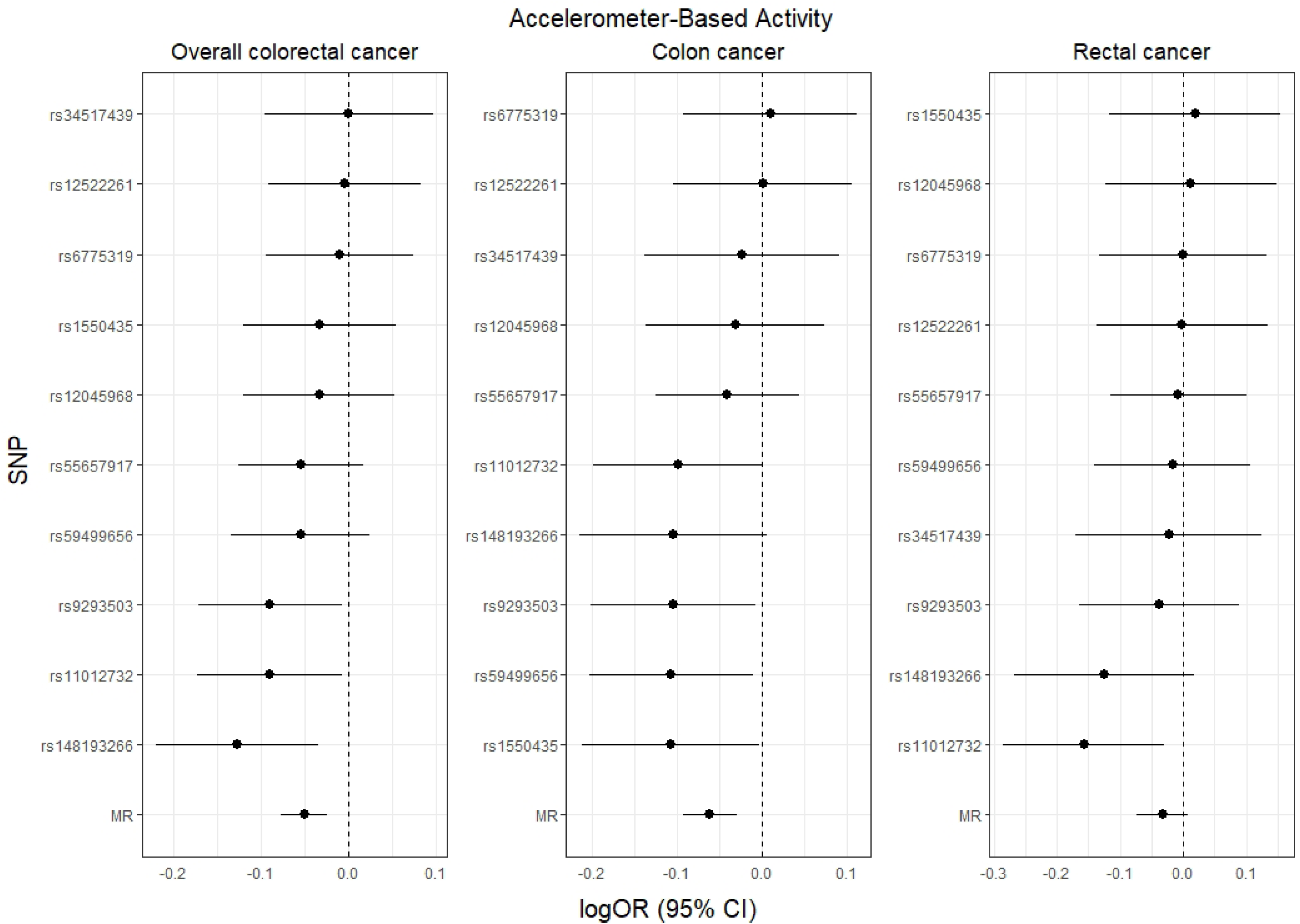
Mendelian randomization analysis for individual SNPs associated with accelerometer-measured physical activity in relation to colorectal cancer risk (overall, colon, rectal). The x axis corresponds to a log OR per one unit increase in the physical activity based on the average acceleration (milli-gravities). The Mendelian randomization (MR) result corresponds to a random effects model due to heterogeneity across the genetic instruments. logOR = log odds ratio. 95% CI = 95% confidence interval. SNP = single nucleotide polymorphism.

Similar results were generally observed for all breast cancer and colorectal cancer endpoints when MR analyses were conducted with the two genome-wide significant accelerometer-measured physical activity SNPs only (Table 2).

### Evaluation of assumptions and sensitivity analyses

The strength of the genetic instruments denoted by the F-statistic was ≥10 for all the accelerometer-measured physical activity variants and ranged between 29 and 56 (Table 1). The intercept test from the MR-Egger regression was statistically significant in the analysis of colorectal cancer in women denoting potential pleiotropy; however, the corrected estimate from MR-Egger replicated the initial finding (Table 2). The estimates from the weighted median approach were consistent with those of inverse variance weighted (IVW) models. The MR pleiotropy residual sum and outlier test (MR-PRESSO) method identified the SNPs rs11012732 and rs55657917 as pleiotropic for breast cancer, but similar magnitude inverse relationships were observed when these variants were excluded from the analyses (Supplementary Table 6).

After examining Phenoscanner and GWAS catalog, we found that several of the accelerometer-measured physical activity genetic variants were also associated with adiposity related phenotypes (Supplementary Table 7). However, the results from the leave–one–SNP out analysis did not reveal any influential SNPs driving the associations (Supplementary Tables 8 – 10). Additionally, similar results were found when the five adiposity-related SNPs were excluded from the genetic instrument (Supplementary Table 11). Further, the results from the multivariable MR analyses adjusting for BMI were largely unchanged from the main IVW results (Supplementary Tables 12, 13).

The association of genetically predicted physical activity and colorectal cancer was similar (OR: 0.60, 95% CI: 0.47 to 0.76, P-value=2.5*10^−5^) after excluding UK Biobank participants from the GWAS for colorectal cancer.

## Discussion

In this MR analysis, higher levels of genetically-predicted accelerometer-measured physical activity were associated with lower risks of breast cancer and colorectal cancer, with similar magnitude inverse associations found for breast cancer subtypes and by colorectal anatomical subsite. These findings indicate that population-level increases in physical activity may lower the incidence of these two commonly diagnosed cancers, and support the promotion of physical activity for cancer prevention.

A large body of observational studies has investigated how physical activity relates to risk of breast and colorectal cancer ^13^. In a participant-level pooled analysis of 12 prospective studies, when the 90^th^ and 10^th^ percentile of leisure-time physical activity were compared, lower risks of breast cancer (Hazard ratio [HR]: 0.90, 95% CI: 0.87 to 0.93), colon cancer (HR: 0.84, 95% CI: 0.77 to 0.91), and rectal cancer (HR: 0.87, 95% CI: 0.80 to 0.95) were found ^3^. These observational studies relied on self-report physical activity assessment methods that are prone to measurement error, which may attenuate associations towards the null. In addition, causality cannot be ascertained from such observational analyses as they are vulnerable to residual confounding and reverse causality. Further, logistical and financial challenges prohibit randomized controlled trials of physical activity and cancer development. For example, it has been estimated that in order to detect a 20% breast cancer risk reduction, between 26,000 to 36,000 healthy middle-aged women would need to be randomized to a 5 year exercise intervention ^14^. Several trials on cancer survivors are registered and underway, and these may provide evidence of potential causal associations between physical activity and disease free survival and cancer recurrence ^15^; however, these interventions will not inform causal inference of the relationship between physical activity and cancer development. We conducted MR analyses to allow causal inference between accelerometer-measured physical activity and risks of developing breast and colorectal cancer. The inverse associations we found were consistent for breast cancer subtypes and across colorectal cancer subsites, and are strongly concordant with prior observational epidemiological evidence.

Being physically active is associated with less weight gain and body fatness, and lower adiposity is associated with lower risks of breast and colorectal cancer ^16,17^. Since body size/adiposity is likely on the causal pathway linking physical activity and breast and colorectal cancer, it is challenging to disentangle independent effects of physical activity on cancer development. This overlap between adiposity and physical activity is evident from 5 of the 10 SNPs in the genetic instrument for accelerometer-measured physical activity previously being associated with adiposity/body size traits. However, it is noteworthy that our results were unchanged when we excluded adiposity-related SNPs from the genetic instrument, and when we conducted multivariable MR analyses adjusting for body mass index (BMI). These results would therefore suggest that physical activity is also associated with breast and colorectal cancer independently of adiposity.

Multiple biological mechanisms are hypothesized to mediate the potential beneficial role of physical activity on cancer development. Greater physical activity has been associated with lower circulating levels of insulin and insulin-like growth factors, which promote cellular proliferation in breast and colorectal tissue and have also been linked to development of cancers at these sites ^18-23^. Higher levels of physical activity have also been associated with lower circulating levels of estradiol, estrone, and higher levels of sex hormone binding globulin ^24-26^ which are strong risk factors for breast cancer development ^27,28^. Physical activity has also been associated with improvements in immune response, with increased surveillance and elimination of cancerous cells ^29,30^. Higher levels of physical activity may also reduce systemic inflammation by lowering the levels of pro-inflammatory factors, such as C-reactive protein (CRP), interleukin-6 (IL-6) and tumour necrosis factor-alpha (TNF-a) ^29,31,32^. Finally, emerging evidence suggests that the gut microbiome may play an important role in the physical activity and cancer relationship. Dysbiosis of the gut microbiome has been associated with increased risks of several malignancies, including breast and colorectal cancer ^33^. Changes in gut microbiome composition and derived metabolic products have been found following endurance exercise training, with short-chain fatty acid concentrations increased in lean, but not obese subjects ^34,35^.

A fundamental assumption of MR is that the genetic variants do not influence the outcome via a different biological pathway from the exposure of interest (horizontal pleiotropy). We conducted multiple sensitivity analyses to test for the influence of pleiotropy on our causal estimates, and our results were robust according to these various tests. A potential limitation of our analysis is that the genetic variants explained a small fraction of the variability of accelerometer-measured physical activity, which may have resulted in some of the breast cancer subtype and colorectal subsite analyses being underpowered. In addition, our use of summary-level data precluded subgroup analyses by other cancer risk factors (e.g. BMI, exogenous hormone use). We were also unable to stratify breast cancer analyses by menopausal status; however, the majority of women in the source GWAS had postmenopausal breast cancer ^11^. Finally, 7-day accelerometer-measured physical activity levels of UK Biobank participants may not have been representative of usual behavioral patterns.

In conclusion, we found that genetically elevated levels of accelerometer-measured physical activity were associated with lower risks of breast and colorectal cancer. These findings strongly support the promotion of physical activity as an effective strategy in the primary prevention of these commonly diagnosed cancers.

## Methods

### Data on physical activity

Summary-level data were obtained from a recently published GWAS on accelerometer-measured physical activity conducted within UK Biobank ^10^. In this GWAS, the regression models were adjusted for age, sex, the first ten genomic principal components, center, season (month), and genotyping chip. This GWAS identified 2 genome-wide-significant polymorphisms (P-value<5×10^−8^) associated with accelerometer-measured physical activity. The estimated SNP-based heritability was 14% suggesting that additional SNPs contributed to its variation. Consequently, for our primary analyses, we used a larger number of 10 independent (linkage disequilibrium [LD] r^2^ ≤ 0.001) genetic variants by relaxing the significance threshold to P-value<1×10^−7^. The expanded number of genetic variants in the accelerometer-measured physical activity instrument also allowed sensitivity analyses to be conducted to check for the influence of horizontal pleiotropy on the results. Data for the associations between the 8 additional SNPs and physical activity were obtained from a recent MR study on physical activity and depression that used the data from the same UK Biobank GWAS ^36^. In secondary analyses, we used the two genome-wide significant SNPs only. Detailed information on the selected genetic variants is provided in Table 1.

### Data on breast cancer and colorectal cancer

Summary data for the associations of the 10 accelerometer-measured genetic variants with breast cancer (overall and by estrogen receptor status: ER positive and ER negative) were obtained from a GWAS of 228,951 women (122,977 breast cancer [69,501 ER positive, 21,468 ER negative] cases and 105,974 controls) of European ancestry from the Breast Cancer Association Consortium (BCAC) ^11^. Genotypes were imputed using the 1000 Genomes Project reference panel and the regression models adjusted for the first ten principal components and country or study (Supplementary Table 1). For colorectal cancer, summary data from 125,915 participants (58,221 colorectal cancer cases and 67,694 controls) were drawn from a meta-analysis within the ColoRectal Transdisciplinary Study (CORECT), the Colon Cancer Family Registry (CCFR), and the Genetics and Epidemiology of Colorectal Cancer (GECCO) consortia ^12^. Imputation was performed using the Haplotype Reference Consortium (HRC) r1.0 reference panel and the regression models were further adjusted for age, sex, genotyping platform (whenever appropriate), and genomic principal components (from 3 to 13, whenever appropriate) (Supplementary Tables 2, 3).

### Statistical power

The a priori statistical power was calculated using an online tool at http://cnsgenomics.com/shiny/mRnd/ ^37^. The 10 accelerometer-measured physical activity SNPs collectively explained 0.4% of phenotypic variability. Given a type 1 error of 5%, we had sufficient power (>80%) when the expected OR per 1 SD was ≤0.83 and ≤0.77 for overall breast cancer (122,977 cases and 105,974 controls) and colorectal cancer (58,221 colorectal cases and 67,694 controls), respectively. The power estimates for subtypes of breast cancer and by subsites of colorectal cancer are presented in Supplementary Table 4.

### Statistical analysis

A two-sample MR approach using summary data and the fixed–effect IVW method was implemented. All accelerometer-measured physical activity and cancer results correspond to an OR per 1 SD increment (8.14 milli-gravities) in the genetically predicted overall average acceleration. The heterogeneity of causal effects by cancer subtype and sex was investigated by estimating the I^2^ statistic assuming a fixed-effects model ^38^.

For causal estimates from MR studies to be valid, three main assumptions must be met: 1) the genetic instrument is strongly associated with the level of accelerometer-measured physical activity; 2) the genetic instrument is not associated with any potential confounder of the physical activity – cancer association; and 3) the genetic instrument does not affect cancer independently of physical activity (i.e. horizontal pleiotropy should not be present) ^39^. The strength of each instrument was measured by calculating the F statistic using the following formula: *F* = *R*^2^ (*N*-2)/(1-*R*^2^), where R^2^ is the proportion of the variability of the physical activity explained by each instrument and N the sample size of the GWAS for the SNP-physical activity association ^40^. To calculate R^2^ we used the following formula: (2 × EAF × (1 - EAF) × beta^2^)/[(2 × EAF × (1-EAF) × beta^2^) + (2 × EAF × (1-EAF) × N× SE(beta)^2^)], where EAF is the effect allele frequency, beta is the estimated genetic effect on physical activity, N is the sample size of the GWAS for the SNP-physical activity association and SE (beta) is the standard error of the genetic effect ^41^.

### Sensitivity analyses

Several sensitivity analyses were used to check and correct for the presence of pleiotropy in the causal estimates. Cochran’s Q was computed to quantify heterogeneity across the individual causal effects, with a P-value≤0.05 indicating the presence of pleiotropy, and that consequently, a random effects IVW MR analysis should be used ^38,42^. We also assessed the potential presence of horizontal pleiotropy using MR-Egger regression based on its intercept term, where deviation from zero denotes the presence of pleiotropy. Additionally, the slope of the MR-Egger regression provides valid MR estimates in the presence of horizontal pleiotropy when the pleiotropic effects of the genetic variants are independent from the genetic associations with the exposure ^43,44^. We also computed OR estimates using the complementary weighted-median method that can give valid MR estimates under the presence of horizontal pleiotropy when up to 50% of the included instruments are invalid ^39^. The presence of pleiotropy was also assessed using the MR-PRESSO. In this, outlying SNPs are excluded from the accelerometer-measured physical activity instrument and the effect estimates are reassessed ^45^. A leave–one–SNP out analysis was also conducted to assess the influence of individual variants on the observed associations. We also examined the selected genetic instruments and their proxies (r^2^>0.8) and their associations with secondary phenotypes (P-value<5×10^−8^) in Phenoscanner (http://www.phenoscanner.medschl.cam.ac.uk/) and GWAS catalog (date checked April 2019).

We also conducted multivariable MR analyses to adjust for potential pleiotropy due to BMI because the initial GWAS on physical activity reported several strong associations (P-value<10^−5^) between the identified SNPs and BMI ^46^. The new estimates correspond to the direct causal effect of physical activity with the BMI being fixed. The genetic data on BMI were obtained from a GWAS study published by The Genetic Investigation of ANthropometric Traits (GIANT) consortium ^47^ (Supplementary Table 5). Additionally, we also conducted analyses with adiposity related SNPs (i.e. those previously associated with BMI, waist circumference, weight, or body/trunk fat percentage in GWAS studies at P-value<10^−8^) excluded (n=5; rs34517439, rs6775319, rs11012732, rs1550435, rs59499656).

Finally, as the GECCO consortium includes 26,763 participants from the UK Biobank, we re-ran the colorectal cancer analyses using GWAS summary estimates with UK Biobank participants excluded in order to correct for any bias and inflated Type 1 errors this may have introduced into our results ^48^.

All the analyses were conducted using the MendelianRandomization package and R programming language ^49^.

## Data availability

Data supporting the findings of this study are available within the paper and its supplementary information files.

## Acknowledgements

This work was supported by a Cancer Research UK program grant (C18281/A19169 to RMM, SJL & NK). RMM was supported by the National Institute for Health Research (NIHR) Bristol Biomedical Research Centre. The views expressed are those of the author(s) and not necessarily those of the NIHR or the Department of Health and Social Care. The funding sources for BCAC, CCFR, GECCO, and CORECT consortia are presented in detail in the appendix in the Supplementary material.

## Author contributions

Study conception: MJG and NM. Data analysis: NP and NM. Drafting of manuscript: NP, MJG, and NM. All other authors contributed to the interpretation of the results and critical revision of the manuscript.

## Competing interests

The authors declare no competing interests

