## Supplementary materials for "Physical activity and risks of breast and colorectal cancer: A Mendelian randomization analysis"

**Supplementary Table 1:** Summary information on breast cancer risk for the 10 SNPs used as genetic instruments for Mendelian randomization analyses

**Supplementary Table 2:** Summary information on colorectal cancer risk for the 10 SNPs used as genetic instruments for Mendelian randomization analyses

**Supplementary Table 3:** Summary information on colorectal cancer risk by subsite for the 10 SNPs used as genetic instruments for Mendelian randomization analyses

**Supplementary Table 4:** Sample size and power calculations in Mendelian randomization study of physical activity and risk of breast and colorectal cancer

**Supplementary Table 5:** Summary information on BMI for 8 SNPs* used as genetic instruments for the multivariable Mendelian randomization analyses

**Supplementary Table 6:** Mendelian randomization estimates between accelerometer-measured physical activity and breast cancer risk, a sensitivity analysis excluding outlying SNPs detected by MR-PRESSO

**Supplementary Table 7:** Evidence of association (p<5*10^-8^) of the 10 SNPs used as genetic instruments for Mendelian randomization analyses of physical activity and risk of breast and colorectal cancer with secondary phenotypes

**Supplementary Table 8:** Mendelian randomization estimates between accelerometer-measured physical activity and breast cancer risk, a sensitivity analysis leaving one SNP out at a time

**Supplementary Table 9:** Mendelian randomization estimates between accelerometer-measured physical activity and colorectal cancer risk overall and by sex, a sensitivity analysis leaving one SNP out at a time

**Supplementary Table 10:** Mendelian randomization estimates between accelerometer-measured physical activity and colorectal cancer risk by subsite, a sensitivity analysis leaving one SNP out at a time

**Supplementary Table 11:** Mendelian randomization estimates between accelerometer-measured physical activity and cancer risk excluding adiposity related SNPs (n=5)

**Supplementary Table 12:** Associations of the genetic instruments with BMI both overall and by sex

**Supplementary Table 13:** Multivariable Mendelian randomization estimates between accelerometer-measured physical activity and cancer risk adjusting for BMI

**Supplementary Figure 1:** Mendelian randomization analysis for individual SNPs associated with accelerometer-measured physical activity in relation to colorectal cancer risk (overall and by anatomical subsite)

**Appendix:** Funding

| **Supplementary Table 1: Summary information on breast cancer risk for the 10 SNPs used as genetic instruments for Mendelian randomization analyses** | | | | | | | | | | | |
| --- | --- | --- | --- | --- | --- | --- | --- | --- | --- | --- | --- |
| **SNP** | **Effect allele** | **Baseline allele** | **Coef. for overall breast cancer** | **SE for overall breast cancer** | **P-value** | **Coef. for ER^+^ breast cancer** | **SE for ER^+^ breast cancer** | **P-value** | **Coef. for ER^-^ breast cancer** | **SE for ER^-^ breast cancer** | **P-value** |
| rs12045968 | G | T | -0.0132 | 0.0078 | 0.09 | -0.0191 | 0.0093 | 0.04 | -0.0193 | 0.0141 | 0.17 |
| rs34517439 | C | A | -0.0104 | 0.0105 | 0.32 | -0.0157 | 0.0126 | 0.21 | -0.0054 | 0.0196 | 0.78 |
| rs6775319 | A | T | -0.0012 | 0.0069 | 0.86 | 0.0034 | 0.0082 | 0.68 | -0.0098 | 0.0126 | 0.44 |
| rs12522261 | G | A | 0.0016 | 0.0066 | 0.81 | -0.0015 | 0.0079 | 0.85 | 0.0036 | 0.0121 | 0.77 |
| rs9293503 | T | C | -0.0102 | 0.0101 | 0.31 | -0.0194 | 0.0121 | 0.11 | -0.0248 | 0.0182 | 0.17 |
| rs11012732 | A | G | -0.0416 | 0.0066 | 2.3×10^-10^ | -0.0586 | 0.0078 | 7.2×10^-14^ | 0.0384 | 0.0121 | 0.002 |
| rs148193266 | C | A | -0.0072 | 0.0186 | 0.7 | -0.0035 | 0.0223 | 0.87 | -0.0228 | 0.034 | 0.5 |
| rs1550435 | T | C | -0.0101 | 0.0063 | 0.11 | -0.0099 | 0.0075 | 0.19 | -0.0096 | 0.0115 | 0.41 |
| rs55657917 | G | T | -0.0483 | 0.0076 | 2.3×10^-10^ | -0.0441 | 0.0091 | 1.4×10^-6^ | -0.0394 | 0.0139 | 0.005 |
| rs59499656 | T | A | -0.0077 | 0.0066 | 0.24 | -0.0116 | 0.0078 | 0.14 | -0.0042 | 0.012 | 0.73 |
| Abbreviations: Coef, coefficient; ER, estrogen receptor; SE, standard error; SNP, single nucleotide polymorphism | | | | | | | | | | | |

| **Supplementary Table 2: Summary information on colorectal cancer risk for the 10 SNPs used as genetic instruments for Mendelian randomization analyses** | | | | | | | | | | | |
| --- | --- | --- | --- | --- | --- | --- | --- | --- | --- | --- | --- |
| **SNP** | **Effect allele** | **Baseline allele** | **Coef. for overall crc cancer** | **SE for overall crc cancer** | **P-value** | **Coef. for crc cancer in men** | **SE for crc cancer in men** | **P-value** | **Coef. for crc cancer in women** | **SE for crc cancer in women** | **P-value** |
| rs12045968 | G | T | -0.008 | 0.0105 | 0.45 | 0.0018 | 0.0146 | 0.9 | -0.0181 | 0.015 | 0.23 |
| rs34517439 | C | A | 0.0001 | 0.0151 | 0.99 | -0.0103 | 0.0211 | 0.63 | 0.0101 | 0.0218 | 0.64 |
| rs6775319 | A | T | -0.0023 | 0.0097 | 0.81 | -0.0184 | 0.0135 | 0.18 | 0.0147 | 0.0139 | 0.29 |
| rs12522261 | G | A | -0.0009 | 0.0094 | 0.92 | 0.0009 | 0.0131 | 0.94 | -0.0036 | 0.0134 | 0.79 |
| rs9293503 | T | C | -0.0294 | 0.0138 | 0.03 | -0.0016 | 0.0192 | 0.93 | -0.0601 | 0.0198 | 0.002 |
| rs11012732 | A | G | -0.0203 | 0.0095 | 0.03 | -0.0176 | 0.0133 | 0.19 | -0.0242 | 0.0137 | 0.08 |
| rs148193266 | C | A | -0.0649 | 0.024 | 0.007 | -0.0331 | 0.0334 | 0.32 | -0.091 | 0.0348 | 0.009 |
| rs1550435 | T | C | -0.0066 | 0.0088 | 0.45 | 0.0003 | 0.0123 | 0.98 | -0.0118 | 0.0127 | 0.35 |
| rs55657917 | G | T | -0.0162 | 0.0109 | 0.13 | 0.0099 | 0.0152 | 0.51 | -0.041 | 0.0156 | 0.008 |
| rs59499656 | T | A | -0.0124 | 0.0092 | 0.18 | -0.01 | 0.0129 | 0.43 | -0.011 | 0.0132 | 0.41 |
| Abbreviations: Coef, coefficient; crc, colorectal; SE, standard error; SNP, single nucleotide polymorphism | | | | | | | | | | | |

| **Supplementary Table 3: Summary information on colorectal cancer risk by subsite for the 10 SNPs used as genetic instruments for Mendelian randomization analyses** | | | | | | | | | | | | | | |
| --- | --- | --- | --- | --- | --- | --- | --- | --- | --- | --- | --- | --- | --- | --- |
| **SNP** | **Effect allele** | **Baseline allele** | **Coef. for colon cancer** | **SE for colon cancer** | **P-value** | **Coef. for proximal colon cancer** | **SE for proximal colon cancer** | **P-value** | **Coef. for distal colon cancer** | **SE for distal colon cancer** | **P-value** | **Coef. for rectal cancer** | **SE for rectal cancer** | **P-value** |
| rs12045968 | G | T | -0.0075 | 0.0128 | 0.55 | -0.0106 | 0.0165 | 0.52 | -0.012 | 0.0168 | 0.48 | 0.003 | 0.0165 | 0.98 |
| rs34517439 | C | A | -0.0074 | 0.018 | 0.68 | -0.0161 | 0.0232 | 0.49 | -0.01 | 0.0238 | 0.67 | -0.007 | 0.0231 | 0.76 |
| rs6775319 | A | T | 0.0022 | 0.0117 | 0.84 | 0.0121 | 0.0151 | 0.42 | -0.0091 | 0.0155 | 0.56 | -0.0002 | 0.0152 | 0.99 |
| rs12522261 | G | A | 0.0002 | 0.0113 | 0.99 | -0.0013 | 0.0145 | 0.93 | -0.0032 | 0.0148 | 0.83 | -0.0004 | 0.0146 | 0.98 |
| rs9293503 | T | C | -0.0342 | 0.0162 | 0.03 | -0.0272 | 0.021 | 0.2 | -0.0407 | 0.0213 | 0.06 | -0.0126 | 0.0213 | 0.55 |
| rs11012732 | A | G | -0.0222 | 0.0114 | 0.05 | -0.0233 | 0.0147 | 0.11 | -0.0207 | 0.015 | 0.16 | -0.0354 | 0.0146 | 0.02 |
| rs148193266 | C | A | -0.0529 | 0.0287 | 0.06 | -0.0557 | 0.0375 | 0.14 | -0.0672 | 0.0383 | 0.08 | -0.0637 | 0.0373 | 0.09 |
| rs1550435 | T | C | -0.0214 | 0.0107 | 0.04 | -0.0093 | 0.0138 | 0.5 | -0.0288 | 0.0141 | 0.04 | 0.0038 | 0.0138 | 0.79 |
| rs55657917 | G | T | -0.0122 | 0.0129 | 0.34 | -0.0025 | 0.0165 | 0.88 | -0.0238 | 0.0169 | 0.16 | -0.0022 | 0.0166 | 0.89 |
| rs59499656 | T | A | -0.0243 | 0.0112 | 0.03 | -0.0163 | 0.0145 | 0.26 | -0.0424 | 0.0147 | 0.004 | -0.0038 | 0.0144 | 0.79 |
| Abbreviations: Coef, coefficient; SE, standard error; SNP, single nucleotide polymorphism | | | | | | | | | | | | | | |

| **Supplementary Table 4: Sample size and power calculations in Mendelian randomization study of physical activity and risk of breast and colorectal cancer** | | | | | | |
| --- | --- | --- | --- | --- | --- | --- |
| **Outcome** | **Sample size** | **Proportion of cases** | **Selected scenarios^*^** | | | |
|  |  |  | **OR=0.90** | **OR=0.85** | **OR=0.80** | **OR=0.75** |
| **Breast cancer** |  |  |  |  |  |  |
| Overall | 228,951 | 0.54 | 0.36 | 0.69 | 0.93 | 0.99 |
| ER^+ve^ | 175,475 | 0.40 | 0.27 | 0.55 | 0.81 | 0.95 |
| ER^-ve^ | 127,442 | 0.17 | 0.14 | 0.26 | 0.42 | 0.60 |
| **Colorectal cancer** |  |  |  |  |  |  |
| Overall | 125,915 | 0.46 | 0.22 | 0.44 | 0.70 | 0.89 |
| Men | 65,815 | 0.47 | 0.14 | 0.26 | 0.44 | 0.64 |
| Women | 59,663 | 0.45 | 0.13 | 0.24 | 0.4 | 0.59 |
| Colon | 98,777 | 0.31 | 0.16 | 0.31 | 0.50 | 0.71 |
| Proximal colon | 81,551 | 0.17 | 0.11 | 0.18 | 0.29 | 0.43 |
| Distal colon | 83,000 | 0.18 | 0.11 | 0.19 | 0.31 | 0.45 |
| Rectal | 83,469 | 0.19 | 0.11 | 0.20 | 0.32 | 0.47 |
| Abbreviations: ER, estrogen receptor; OR, odds ratio  ^*^ Type 1 error of 5% and a proportion of variance explained equal to 0.4% are assumed | | | | | | |

| **Supplementary Table 5: Summary information on BMI for 8 SNPs^*^ used as genetic instruments for the multivariable Mendelian randomization analyses** | | | | | | | | |
| --- | --- | --- | --- | --- | --- | --- | --- | --- |
| **SNP** | **Effect allele** | **Baseline allele** | **Coef. BMI overall** | **SE BMI overall** | **Coef. BMI men** | **SE BMI men** | **Coef. BMI women** | **SE BMI women** |
| rs12045968 | G | T | 0.0065 | 0.0044 | 0.0054 | 0.006 | 0.0088 | 0.0055 |
| rs6775319^†^ | A | T | -0.0089 | 0.0041 | -0.0071 | 0.0055 | -0.01 | 0.0052 |
| rs12522261 | G | A | 0.0078 | 0.004 | 0.0023 | 0.0053 | 0.0126 | 0.005 |
| rs9293503^†^ | T | C | 0.0057 | 0.0063 | 0.0014 | 0.0085 | 0.0094 | 0.008 |
| rs11012732^†^ | A | G | -0.0127 | 0.0042 | -0.0085 | 0.0057 | -0.0156 | 0.0052 |
| rs1550435 | T | C | -0.003 | 0.0038 | -0.001 | 0.0052 | -0.0061 | 0.0049 |
| rs55657917^†^ | G | T | 0.0034 | 0.0047 | -0.0018 | 0.0062 | 0.0075 | 0.0059 |
| rs59499656^†^ | T | A | -0.012 | 0.004 | -0.0028 | 0.0053 | -0.0209 | 0.005 |
| Abbreviations: BMI, body mass index; Coef, coefficient; SE; standard error  ^*^ rs148193266 and rs34517439 were excluded since they were unavailable in the BMI GWAS and no good proxies (r^2^>0.8) were found  ^†^ These SNP were unavailable in BMI GWAS and proxies were used instead | | | | | | | | |

| **Supplementary Table 6: Mendelian randomization estimates between accelerometer-measured physical activity and breast cancer risk, a sensitivity analysis excluding outlying SNPs detected by MR-PRESSO** | | | | | | | | | | | | |
| --- | --- | --- | --- | --- | --- | --- | --- | --- | --- | --- | --- | --- |
|  | **Overall breast cancer**  **(exclude rs11012732, rs55657917)** | | | | **ER^+ve^ subset**  **(exclude rs11012732)** | | | | **ER^-ve^ subset**  **(exclude rs11012732)** | | | |
|  | **OR** | **95% CI** | **P-value** | **P-value for pleiotropy or heterogeneity** | **OR** | **95% CI** | **P-value** | **P-value for pleiotropy or heterogeneity** | **OR** | **95% CI** | **P-value** | **P-value for pleiotropy or heterogeneity** |
| Inverse-variance weighted | 0.80 | 0.67, 0.96 | 0.02 | 0.85 | 0.64 | 0.48, 0.85 | 0.002 | 0.04 | 0.64 | 0.48, 0.87 | 0.004 | 0.7 |
| MR-Egger | 0.83 | 0.37, 1.84 | 0.65 | 0.94 | 0.42 | 0.11, 1.58 | 0.20 | 0.51 | 0.34 | 0.09, 1.35 | 0.13 | 0.36 |
| Weighted median | 0.76 | 0.61, 0.96 | 0.02 |  | 0.66 | 0.50, 0.88 | 0.004 |  | 0.69 | 0.47, 1.02 | 0.07 |  |
| Abbreviations: ER, estrogen receptor; OR, odds ratio | | | | | | | | | | | | |

| **Supplementary Table 7: Evidence of association (p<5*10^-8^) of the 10 SNPs used as genetic instruments for Mendelian randomization analyses of physical activity and risk of breast and colorectal cancer with secondary phenotypes** | | | |
| --- | --- | --- | --- |
| **SNP** | **Chr** | **Gene** | **Diseases & traits** |
| rs12045968 | 1 | ZNF362 | None found |
| rs34517439 | 1 | DNAJB4 | Lung cancer (PMID: 28604730), Diastolic blood pressure (PMID: 30224653)  Height (UK biobank), Basal metabolic rate (UK biobank), Weight (UK biobank)  Hip circumference (UK biobank), Waist circumference (UK biobank)  ΒΜΙ (UK biobank), Creatinine in urine (UK biobank)  Psoriasis (PMID: 28537254) |
| rs6775319 | 3 | LOC105376976 | Body/trunk fat percentage (UK biobank) |
| rs12522261 | 5 | LINC01470 | Getting up in morning (UK Biobank)  Morning or evening person (UK Biobank, SNP in LD: rs12517065) |
| rs9293503 | 5 | LINC00461 | None found |
| rs11012732 | 10 | MLLT10 | Meningioma (PMID: 21804547), Ovarian cancer(PMID: 28346442)  Breast cancer (PMID:29059683, SNP in LD: rs7098100), Waist circumference (UK biobank)  Hip circumference (UK biobank), ΒΜΙ (UK biobank), Weight (UK biobank)  Sodium in urine (UK biobank), Creatinine in urine (UK biobank SNP in LD: rs61850044) |
| rs148193266 | 11 | RP11-681H10.1 | None found |
| rs1550435 | 15 | PML | Height (UK biobank and other GWAS), ΒΜΙ (UK biobank, SNP in LD: rs9479) |
| rs55657917 | 17 | CRHR1 | Height (UK biobank)  Alcohol intake frequency (UK biobank)  Systolic blood pressure (UK biobank)  Parkinson’s disease (PMID: 21292315, SNP in LD: rs2942168)  Red blood cell count (PMID: 27863252)  Ovarian cancer(PMID: 28346442) |
| rs59499656 | 18 | RIT2/SYT4 | ΒΜΙ (UK biobank)  Weight (UK biobank) |
| Abbreviations: Chr, chromosome; PMID, Pubmed ID; SNP, single nucleotide polymorphism | | | |

| **Supplementary Table 8: Mendelian randomization estimates^*^ between accelerometer-measured physical activity and breast cancer risk, a sensitivity analysis leaving one SNP out at a time** | | | | | | | | | | | | | | | |
| --- | --- | --- | --- | --- | --- | --- | --- | --- | --- | --- | --- | --- | --- | --- | --- |
| **Excluded SNP** | **Overall breast cancer** | | | | | **ER+ve subset** | | | | | **ER-ve subset** | | | | |
|  | **OR** | **95% CI** | **P-value** | **P-value het**^†^ | **P-value inter**^‡^ | **OR** | **95% CI** | **P-value** | **P-value het**^†^ | **P-value inter**^‡^ | **OR** | **95% CI** | **P-value** | **P-value het**^†^ | **P-value inter**^‡^ |
| rs12045968 | 0.59 | 0.40, 0.87 | 7.6×10^-3^ | 2.8×10^-7^ | 0.94 | 0.53 | 0.33, 0.85 | 8.9×10^-3^ | 1.2×10^-7^ | 0.91 | 0.82 | 0.51, 1.31 | 0.41 | 0.01 | 0.23 |
| rs34517439 | 0.58 | 0.40, 0.85 | 5.4×10^-3^ | 3.9×10^-7^ | 0.86 | 0.52 | 0.33, 0.83 | 6.4×10^-3^ | 1.5×10^-7^ | 0.96 | 0.78 | 0.48, 1.26 | 0.31 | 0.01 | 0.24 |
| rs6775319 | 0.56 | 0.39, 0.82 | 2.5×10^-3^ | 1.6×10^-6^ | 0.93 | 0.49 | 0.32, 0.76 | 1.3×10^-3^ | 2.8×10^-6^ | 0.71 | 0.80 | 0.49, 1.29 | 0.35 | 0.01 | 0.24 |
| rs12522261 | 0.56 | 0.39, 0.80 | 1.7×10^-3^ | 3.3×10^-6^ | 0.81 | 0.5 | 0.32, 0.79 | 2.8×10^-3^ | 6.7×10^-7^ | 0.69 | 0.75 | 0.47, 1.21 | 0.25 | 0.01 | 0.33 |
| rs9293503 | 0.58 | 0.39, 0.85 | 5.0×10^-3^ | 4.7×10^-7^ | 0.79 | 0.52 | 0.33, 0.84 | 7.2×10^-3^ | 1.4×10^-7^ | 0.96 | 0.82 | 0.51, 1.32 | 0.41 | 0.01 | 0.33 |
| rs11012732 | 0.67 | 0.50, 0.90 | 8.8×10^-3^ | 8.2×10^-4^ | 0.52 | 0.64 | 0.49, 0.85 | 1.8×10^-3^ | 0.04 | 0.52 | 0.64 | 0.48, 0.87 | 0.004 | 0.70 | 0.36 |
| rs148193266 | 0.58 | 0.40, 0.84 | 4.0×10^-3^ | 6.5×10^-7^ | 0.54 | 0.51 | 0.32, 0.80 | 3.5×10^-3^ | 4.1×10^-7^ | 0.68 | 0.79 | 0.49, 1.28 | 0.34 | 0.01 | 0.17 |
| rs1550435 | 0.59 | 0.40, 0.87 | 7.2×10^-3^ | 3.0×10^-7^ | 0.99 | 0.52 | 0.32, 0.83 | 6.3×10^-3^ | 1.6×10^-7^ | 0.79 | 0.80 | 0.49, 1.29 | 0.36 | 0.01 | 0.17 |
| rs55657917 | 0.68 | 0.50, 0.93 | 1.7×10^-3^ | 3.9×10^-4^ | 0.61 | 0.59 | 0.37, 0.92 | 2.1×10^-2^ | 1.8×10^-6^ | 0.66 | 0.91 | 0.59, 1.39 | 0.65 | 0.05 | 0.43 |
| rs59499656 | 0.58 | 0.39, 0.85 | 5.3×10^-3^ | 4.6×10^-7^ | 1 | 0.52 | 0.32, 0.83 | 6.5×10^-3^ | 1.6×10^-7^ | 0.86 | 0.78 | 0.48, 1.26 | 0.31 | 0.01 | 0.27 |
| Abbreviations: CI, confidence interval; ER, estrogen receptor; het, heterogeneity; inter, intercept; OR, odds ratio; SNP, single nucleotide polymorphism  ^*^ The estimates are derived from a random effects Mendelian Randomization analysis due to the large heterogeneity based on Cochran’s Q test  ^†^ P – value of Cochran’s Q test  ^‡^ P – value of the intercept term from the MR-Egger’s regression | | | | | | | | | | | | | | | |

| **Supplementary Table 9: Mendelian randomization estimates between accelerometer-measured physical activity and colorectal cancer risk overall and by sex, a sensitivity analysis leaving one SNP out at a time** | | | | | | | | | |
| --- | --- | --- | --- | --- | --- | --- | --- | --- | --- |
| **Excluded SNP** | **Both sexes** | | | **Men** | | | **Women** | | |
|  | **OR** | **95% CI** | **P-value** | **OR** | **95% CI** | **P-value** | **OR** | **95% CI** | **P-value** |
| rs12045968 | 0.65 | 0.52, 0.82 | 2.4×10^-4^ | 0.8 | 0.58, 1.10 | 0.18 | 0.55 | 0.39, 0.76 | 2.7×10^-4^ |
| rs34517439 | 0.64 | 0.51, 0.80 | 1.0×10^-4^ | 0.83 | 0.61, 1.14 | 0.25 | 0.51 | 0.37, 0.70 | 3.7×10^-5^ |
| rs6775319 | 0.64 | 0.51, 0.80 | 1.2×10^-4^ | 0.87 | 0.63, 1.19 | 0.39 | 0.48 | 0.35, 0.67 | 1.1×10^-5^ |
| rs12522261 | 0.64 | 0.51, 0.80 | 1.0×10^-4^ | 0.81 | 0.59, 1.11 | 0.18 | 0.52 | 0.37, 0.72 | 8.0×10^-5^ |
| rs9293503 | 0.69 | 0.55, 0.86 | 0.001 | 0.81 | 0.59, 1.11 | 0.2 | 0.6 | 0.43, 0.84 | 0.003 |
| rs11012732 | 0.69 | 0.55, 0.86 | 0.001 | 0.87 | 0.63, 1.19 | 0.38 | 0.56 | 0.40, 0.78 | 0.001 |
| rs148193266 | 0.7 | 0.56, 0.88 | 0.002 | 0.85 | 0.62, 1.17 | 0.31 | 0.59 | 0.42, 0.81 | 0.001 |
| rs1550435 | 0.65 | 0.52, 0.82 | 2.4×10^-4^ | 0.81 | 0.59, 1.11 | 0.19 | 0.54 | 0.39, 0.74 | 1.9×10^-4^ |
| rs55657917 | 0.67 | 0.53, 0.84 | 0.001 | 0.77 | 0.55, 1.06 | 0.11 | 0.59 | 0.42, 0.83 | 0.002 |
| rs59499656 | 0.67 | 0.53, 0.84 | 0.001 | 0.84 | 0.61, 1.16 | 0.29 | 0.53 | 0.38, 0.74 | 1.6×10^-4^ |
| Abbreviations: CI, confidence interval; OR, odds ratio; | | | | | | | | | |

| **Supplementary Table 10: Mendelian randomization estimates between accelerometer-measured physical activity and colorectal cancer risk by subsite, a sensitivity analysis leaving one SNP out at a time** | | | | | | | | | | | | |
| --- | --- | --- | --- | --- | --- | --- | --- | --- | --- | --- | --- | --- |
| **Excluded SNP** | **Colon cancer** | | | **Proximal colon** | | | **Distal colon** | | | **Rectal cancer** | | |
|  | **OR** | **95% CI** | **P-value** | **OR** | **95% CI** | **P-value** | **OR** | **95% CI** | **P-value** | **OR** | **95% CI** | **P-value** |
| rs12045968 | 0.59 | 0.45, 0.78 | 1.6×10^-4^ | 0.65 | 0.46, 0.93 | 0.02 | 0.65 | 0.46, 0.93 | 0.02 | 0.74 | 0.52, 1.05 | 0.09 |
| rs34517439 | 0.59 | 0.45, 0.78 | 1.4×10^-4^ | 0.64 | 0.45, 0.91 | 0.01 | 0.64 | 0.45, 0.91 | 0.01 | 0.76 | 0.54, 1.08 | 0.12 |
| rs6775319 | 0.57 | 0.43, 0.75 | 5.6×10^-5^ | 0.64 | 0.45, 0.91 | 0.01 | 0.64 | 0.45, 0.92 | 0.01 | 0.74 | 0.52, 1.06 | 0.10 |
| rs12522261 | 0.58 | 0.44, 0.76 | 7.5×10^-5^ | 0.64 | 0.45, 0.91 | 0.01 | 0.64 | 0.44, 0.91 | 0.01 | 0.75 | 0.52, 1.06 | 0.10 |
| rs9293503 | 0.63 | 0.48, 0.83 | 0.001 | 0.69 | 0.48, 0.98 | 0.04 | 0.69 | 0.48, 0.99 | 0.04 | 0.77 | 0.54, 1.10 | 0.15 |
| rs11012732 | 0.63 | 0.48, 0.83 | 0.001 | 0.69 | 0.48, 0.98 | 0.04 | 0.69 | 0.48, 0.98 | 0.04 | 0.86 | 0.60, 1.23 | 0.41 |
| rs148193266 | 0.63 | 0.48, 0.82 | 0.001 | 0.70 | 0.49, 0.99 | 0.05 | 0.70 | 0.49, 1.00 | 0.05 | 0.82 | 0.58, 1.16 | 0.26 |
| rs1550435 | 0.63 | 0.48, 0.83 | 0.001 | 0.65 | 0.46, 0.93 | 0.02 | 0.65 | 0.46, 0.93 | 0.02 | 0.73 | 0.52, 1.04 | 0.08 |
| rs55657917 | 0.59 | 0.45, 0.78 | 2.3×10^-4^ | 0.67 | 0.46, 0.96 | 0.03 | 0.66 | 0.46, 0.96 | 0.03 | 0.74 | 0.51, 1.06 | 0.10 |
| rs59499656 | 0.64 | 0.48, 0.84 | 0.001 | 0.66 | 0.47, 0.95 | 0.02 | 0.66 | 0.46, 0.95 | 0.03 | 0.75 | 0.53, 1.07 | 0.12 |
| Abbreviations: CI, confidence interval; OR, odds ratio; | | | | | | | | | | | | |

| **Supplementary Table 11: Mendelian randomization estimates between accelerometer-measured physical activity and cancer risk excluding adiposity related SNPs (n=5)** | | | | |
| --- | --- | --- | --- | --- |
| **Methods** |  | | | |
|  | **Estimates (OR)^*^** | **95% CI** | **P-value** | **P-value for pleiotropy**^†^ **or heterogeneity**^‡^ |
| **Overall Breast Cancer** | | | | |
| Inverse-variance weighted^§^ | 0.60 | 0.35, 1.01 | 0.053 | 1.0×10^-4^ |
| MR-Egger | 0.43 | 0.04, 5.05 | 0.50 | 0.8 |
| Weighted median | 0.72 | 0.51, 1.02 | 0.07 |  |
| **ER^+ve^ subset** | | | | |
| Inverse-variance weighted^§^ | 0.56 | 0.36, 0.87 | 0.01 | 0.02 |
| MR-Egger | 0.50 | 0.06, 4.19 | 0.53 | 0.90 |
| Weighted median | 0.58 | 0.39, 0.86 | 0.008 |  |
| **ER^-ve^ subset** | | | | |
| Inverse-variance weighted | 0.56 | 0.38, 0.84 | 0.005 | 0.37 |
| MR-Egger | 0.29 | 0.05, 1.81 | 0.19 | 0.47 |
| Weighted median | 0.53 | 0.31, 0.91 | 0.02 |  |
| **Overall Colorectal Cancer** | | | | |
| Inverse-variance weighted | 0.61 | 0.45, 0.83 | 0.001 | 0.34 |
| MR-Egger | 0.18 | 0.05, 0.59 | 0.005 | 0.04 |
| Weighted median | 0.64 | 0.43, 0.98 | 0.04 |  |
| **Colorectal Cancer in men** | | | | |
| Inverse-variance weighted | 1.00 | 0.65, 1.51 | 0.99 | 0.83 |
| MR-Egger | 0.55 | 0.10, 2.93 | 0.48 | 0.47 |
| Weighted median | 1.04 | 0.62, 1.74 | 0.87 |  |
| **Colorectal Cancer in women** | | | | |
| Inverse-variance weighted | 0.38 | 0.24, 0.58 | 1.04×10^-5^ | 0.08 |
| MR-Egger | 0.06 | 0.01, 0.36 | 0.002 | 0.04 |
| Weighted median | 0.33 | 0.18, 0.61 | 3.9×10^-4^ |  |
| **Colon Cancer** | | | | |
| Inverse-variance weighted | 0.61 | 0.45, 0.91 | 0.015 | 0.53 |
| MR-Egger | 0.21 | 0.05, 0.89 | 0.034 | 0.12 |
| Weighted median | 0.55 | 0.39, 0.78 | 0.001 |  |
| **Proximal Colon Cancer** | | | | |
| Inverse-variance weighted | 0.69 | 0.43, 1.09 | 0.12 | 0.76 |
| MR-Egger | 0.27 | 0.04, 1.75 | 0.17 | 0.31 |
| Weighted median | 0.74 | 0.42, 1.32 | 0.31 |  |
| **Distal Colon Cancer** | | | | |
| Inverse-variance weighted | 0.52 | 0.33, 0.84 | 0.007 | 0.74 |
| MR-Egger | 0.15 | 0.02, 1.01 | 0.051 | 0.19 |
| Weighted median | 0.52 | 0.33, 0.84 | 0.007 |  |
| **Rectal Cancer** | | | | |
| Inverse-variance weighted | 0.80 | 0.50, 1.27 | 0.34 | 0.66 |
| MR-Egger | 0.23 | 0.04, 1.51 | 0.13 | 0.18 |
| Weighted median | 0.94 | 0.53, 1.67 | 0.83 |  |
| Abbreviations:CI, confidence intervals; MR: Mendelian Randomization; OR: odds ratio; SNPs: Single nucleotide polymorphisms  ^*^ The estimates correspond to a standard deviation increase in physical activity  ^†^ P-value or pleiotropy based on MR-Egger intercept  ^‡^ P-value for heterogeneity based on Q statistic  ^§^ The estimates were derived from a random-effects model due to the presence of heterogeneity based on Cochran’s Q statistic | | | | |

| **Supplementary Table 12: Associations of the genetic instruments with BMI both overall and by sex** | | | |
| --- | --- | --- | --- |
| **SNP** | **Beta_Overall (se)** | **Beta_Men (se)** | **Beta_Women (se)** |
| rs12045968 | 0.0065 (0.0044) | 0.0054 (0.006) | 0.0088 (0.0055) |
| rs34517439^*^ |  |  |  |
| rs4390955 (proxy for rs6775319) | -0.0089 (0.0041) | -0.0071 (0.0055) | -0.01 (0.0052) |
| rs12522261 | 0.0078 (0.004) | 0.0023 (0.0053) | 0.0126 (0.005) |
| rs10067451 (proxy for rs9293503) | 0.0057 (0.0063) | 0.0014 (0.0085) | 0.0094 (0.008) |
| rs7084454 (proxy for rs11012732) | -0.0127 (0.0042) | -0.0085 (0.0057) | -0.0156 (0.0052) |
| rs148193266^*^ |  |  |  |
| rs1550435 | -0.003 (0.0038) | -0.001 (0.0052) | -0.0061 (0.0049) |
| rs11079724 (proxy for rs55657917) | 0.0034 (0.0047) | -0.0018 (0.0062) | 0.0075 (0.0059) |
| rs2052607 (proxy for rs59499656) | -0.012 (0.004) | -0.0028 (0.0053) | -0.0209 (0.005) |
| ^*^ rs148193266 and rs34517439 were not available in the BMI GWAS and no good proxies could be found (r^2^<0.8) | | | |

| **Supplementary Table 13: Multivariable Mendelian randomization estimates between accelerometer-measured physical activity and cancer risk adjusting for BMI** | | | |
| --- | --- | --- | --- |
| **Cancer type** | **Estimates (OR)^*^** | **95% CI** | **P-value** |
| **Overall Breast Cancer** |  |  |  |
| Inverse-variance weighted^†^ | 0.57 | 0.36, 0.91 | 0.02 |
| **ER+ subset** |  |  |  |
| Inverse-variance weighted^†^ | 0.51 | 0.30, 0.89 | 0.02 |
| **ER- subset** |  |  |  |
| Inverse-variance weighted^†^ | 0.73 | 0.44, 1.22 | 0.22 |
| **Overall Colorectal Cancer** |  |  |  |
| Inverse-variance weighted | 0.69 | 0.54, 0.88 | 0.003 |
| **Colorectal Cancer in men** |  |  |  |
| Inverse-variance weighted | 0.95 | 0.67, 1.35 | 0.78 |
| **Colorectal Cancer in women** |  |  |  |
| Inverse-variance weighted | 0.53 | 0.34, 0.83 | 0.005 |
| **Colon Cancer** |  |  |  |
| Inverse-variance weighted | 0.63 | 0.47, 0.85 | 0.002 |
| **Proximal Colon Cancer** |  |  |  |
| Inverse-variance weighted | 0.75 | 0.51, 1.08 | 0.13 |
| **Distal Colon Cancer** |  |  |  |
| Inverse-variance weighted | 0.50 | 0.34, 0.73 | 3.0×10^-4^ |
| **Rectal Cancer** |  |  |  |
| Inverse-variance weighted | 0.86 | 0.59, 1.27 | 0.45 |
| Abbreviations: BMI, body mass index CI, confidence intervals; OR: odds ratio; SNPs: Single nucleotide polymorphism  ^*^ The estimates correspond to a standard deviation increase in physical activity  ^†^ The estimates were derived from a random-effects model due to the presence of heterogeneity based on Cochran’s Q statistic | | | |

**Supplementary Figure 1: Mendelian randomization analysis for individual SNPs associated with accelerometer-measured physical activity in relation to colorectal cancer risk (overall and by anatomical subsite).** The x axis corresponds to a log OR per one unit increase in the physical activity based on the average acceleration (milli-gravities).

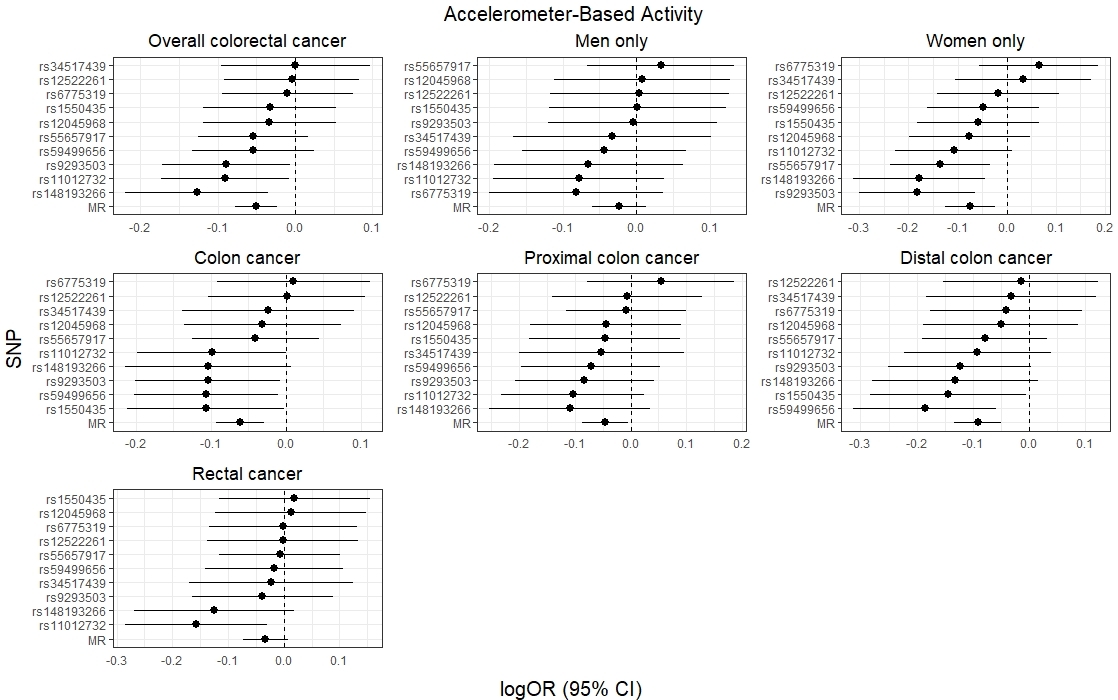

**Appendix:** **Funding**

The breast cancer genome-wide association analyses were supported by the Government of Canada through Genome Canada and the Canadian Institutes of Health Research, the ‘Ministère de l’Économie, de la Science et de l’Innovation du Québec’ through Genome Québec and grant PSR-SIIRI-701, The National Institutes of Health (U19 CA148065, X01HG007492), Cancer Research UK (C1287/A10118, C1287/A16563, C1287/A10710) and The European Union (HEALTH-F2-2009-223175 and H2020 633784 and 634935). All studies and funders are listed in Michailidou et al (2017).

Genetics and Epidemiology of Colorectal Cancer Consortium (GECCO): National Cancer Institute, National Institutes of Health, U.S. Department of Health and Human Services (U01 CA137088, R01 CA059045, R01201407). Genotyping/Sequencing services were provided by the Center for Inherited Disease Research (CIDR). CIDR is fully funded through a federal contract from the National Institutes of Health to The Johns Hopkins University, contract number HHSN268201200008I. This research was funded in part through the NIH/NCI Cancer Center Support Grant P30 CA015704.

ASTERISK: a Hospital Clinical Research Program (PHRC-BRD09/C) from the University Hospital Center of Nantes (CHU de Nantes) and supported by the Regional Council of Pays de la Loire, the Groupement des Entreprises Françaises dans la Lutte contre le Cancer (GEFLUC), the Association Anne de Bretagne Génétique and the Ligue Régionale Contre le Cancer (LRCC).

The ATBC Study is supported by the Intramural Research Program of the U.S. National Cancer Institute, National Institutes of Health, and by U.S. Public Health Service contract HHSN261201500005C from the National Cancer Institute, Department of Health and Human Services.

CLUE II: This research was funded by the American Institute for Cancer Research and the Maryland Cigarette Restitution Fund at Johns Hopkins, and the NCI (P30 CA006973 to W.G. Nelson).

COLO2&3: National Institutes of Health (R01 CA60987).

ColoCare: This work was supported by the National Institutes of Health (grant numbers R01 CA189184 (Li/Ulrich), U01 CA206110 (Ulrich/Li/Siegel/Figueireido/Colditz, 2P30CA015704- 40 (Gilliland), R01 CA207371 (Ulrich/Li)), the Matthias Lackas-Foundation, the German Consortium for Translational Cancer Research, and the EU TRANSCAN initiative.

The Colon Cancer Family Registry (CFR) Illumina GWAS was supported by funding from the National Cancer Institute, National Institutes of Health (grant numbers U01 CA122839, R01 CA143247). The Colon CFR/CORECT Affymetrix Axiom GWAS and OncoArray GWAS were supported by funding from National Cancer Institute, National Institutes of Health (grant number U19 CA148107 to S Gruber). The Colon CFR participant recruitment and collection of data and biospecimens used in this study were supported by the National Cancer Institute, National Institutes of Health (grant number U01 CA167551) and through cooperative agreements with the following Colon CFR centers: Australasian Colorectal Cancer Family Registry (NCI/NIH grant numbers U01 CA074778 and U01/U24 CA097735), USC Consortium Colorectal Cancer Family Registry (NCI/NIH grant numbers U01/U24 CA074799), Mayo Clinic Cooperative Family Registry for Colon Cancer Studies (NCI/NIH grant number U01/U24 CA074800), Ontario Familial Colorectal Cancer Registry (NCI/NIH grant number U01/U24 CA074783), Seattle Colorectal Cancer Family Registry (NCI/NIH grant number U01/U24 CA074794), and University of Hawaii Colorectal Cancer Family Registry (NCI/NIH grant number U01/U24 CA074806), Additional support for case ascertainment was provided from the Surveillance,

Epidemiology and End Results (SEER) Program of the National Cancer Institute to Fred Hutchinson Cancer Research Center (Control Nos. N01-CN-67009 and N01-PC-35142, and Contract No. HHSN2612013000121), the Hawai‘i Department of Health (Control Nos. N01-PC- 67001 and N01-PC-35137, and Contract No. HHSN26120100037C, and the California Department of Public Health (contracts HHSN261201000035C awarded to the University of Southern California, and the following state cancer registries: AZ, CO, MN, NC, NH, and by the Victoria Cancer Registry and Ontario Cancer Registry.

COLON: The COLON study is sponsored by Wereld Kanker Onderzoek Fonds, including funds from grant 2014/1179 as part of the World Cancer Research Fund International Regular Grant Programme, by Alpe d’Huzes and the Dutch Cancer Society (UM 2012–5653, UW 2013-5927, UW2015-7946), and by TRANSCAN (JTC2012-MetaboCCC, JTC2013-FOCUS).

Colorectal Cancer Transdisciplinary (CORECT) Study: The CORECT Study was supported by the National Cancer Institute, National Institutes of Health (NCI/NIH), U.S. Department of Health and Human Services (grant numbers U19 CA148107, R01 CA81488, P30 CA014089, R01 CA197350,; P01 CA196569; R01 CA201407) and National Institutes of Environmental Health Sciences, National Institutes of Health (grant number T32 ES013678).

CORSA: “Österreichische Nationalbank Jubiläumsfondsprojekt” (12511) and Austrian Research Funding Agency (FFG) grant 829675.

CPS-II: The American Cancer Society funds the creation, maintenance, and updating of the Cancer Prevention Study-II (CPS-II) cohort. This study was conducted with Institutional Review Board approval.

CRCGEN: Colorectal Cancer Genetics & Genomics, Spanish study was supported by Instituto de Salud Carlos III, co-funded by FEDER funds –a way to build Europe– (grants PI14-613 and PI09-1286), Agency for Management of University and Research Grants (AGAUR) of the Catalan Government (grant 2017SGR723), and Junta de Castilla y León (grant LE22A10-2).

Sample collection of this work was supported by the Xarxa de Bancs de Tumors de Catalunya sponsored by Pla Director d’Oncología de Catalunya (XBTC), Plataforma Biobancos PT13/0010/0013 and ICOBIOBANC, sponsored by the Catalan Institute of Oncology.

Czech Republic CCS: This work was supported by the Grant Agency of the Czech Republic (grants CZ GA CR: GAP304/10/1286 and 1585) and by the Grant Agency of the Ministry of Health of the Czech Republic (grants AZV 15-27580A and AZV 17-30920A).

DACHS: This work was supported by the German Research Council (BR 1704/6-1, BR 1704/6-

3, BR 1704/6-4, CH 117/1-1, HO 5117/2-1, HE 5998/2-1, KL 2354/3-1, RO 2270/8-1 and BR

1704/17-1), the Interdisciplinary Research Program of the National Center for Tumor Diseases (NCT), Germany, and the German Federal Ministry of Education and Research (01KH0404, 01ER0814, 01ER0815, 01ER1505A and 01ER1505B).

DALS: National Institutes of Health (R01 CA48998 to M. L. Slattery).

EDRN: This work is funded and supported by the NCI, EDRN Grant (U01 CA 84968-06).

EPIC: The coordination of EPIC is financially supported by the European Commission (DGSANCO) and the International Agency for Research on Cancer. The national cohorts are supported by Danish Cancer Society (Denmark); Ligue Contre le Cancer, Institut Gustave Roussy, Mutuelle Générale de l’Education Nationale, Institut National de la Santé et de la Recherche Médicale (INSERM) (France); German Cancer Aid, German Cancer Research Center (DKFZ), Federal Ministry of Education and Research (BMBF), Deutsche Krebshilfe, Deutsches Krebsforschungszentrum and Federal Ministry of Education and Research (Germany); the Hellenic Health Foundation (Greece); Associazione Italiana per la Ricerca sul Cancro-AIRCItaly and National Research Council (Italy); Dutch Ministry of Public Health, Welfare and Sports (VWS), Netherlands Cancer Registry (NKR), LK Research Funds, Dutch Prevention Funds, Dutch ZON (Zorg Onderzoek Nederland), World Cancer Research Fund (WCRF), Statistics Netherlands (The Netherlands); ERC-2009-AdG 232997 and Nordforsk, Nordic Centre of Excellence programme on Food, Nutrition and Health (Norway); Health Research Fund (FIS), PI13/00061 to Granada, PI13/01162 to EPIC-Murcia, Regional Governments of Andalucía, Asturias, Basque Country, Murcia and Navarra, ISCIII RETIC (RD06/0020) (Spain); Swedish Cancer Society, Swedish Research Council and County Councils of Skåne and Västerbotten (Sweden); Cancer Research UK (14136 to EPIC-Norfolk; C570/A16491 and C8221/A19170 to EPIC-Oxford), Medical Research Council (1000143 to EPIC-Norfolk, MR/M012190/1 to EPICOxford) (United Kingdom).

EPICOLON: This work was supported by grants from Fondo de Investigación Sanitaria/FEDER (PI08/0024, PI08/1276, PS09/02368, P111/00219, PI11/00681, PI14/00173, PI14/00230,

PI17/00509, 17/00878, Acción Transversal de Cáncer), Xunta de Galicia (PGIDIT07PXIB9101209PR), Ministerio de Economia y Competitividad (SAF07-64873, SAF 2010-19273, SAF2014-54453R), Fundación Científica de la Asociación Española contra el Cáncer (GCB13131592CAST), Beca Grupo de Trabajo “Oncología” AEG (Asociación Española de Gastroenterología), Fundación Privada Olga Torres, FP7 CHIBCHA Consortium, Agència de Gestió d’Ajuts Universitaris i de Recerca (AGAUR, Generalitat de Catalunya, 2014SGR135, 2014SGR255, 2017SGR21, 2017SGR653), Catalan Tumour Bank Network (Pla Director d’Oncologia, Generalitat de Catalunya), PERIS (SLT002/16/00398, Generalitat de Catalunya), CERCA Programme (Generalitat de Catalunya) and COST Action CA17118. CIBERehd is funded by the Instituto de Salud Carlos III.

ESTHER/VERDI. This work was supported by grants from the Baden-Württemberg Ministry of Science, Research and Arts and the German Cancer Aid.

Harvard cohorts (HPFS, NHS, PHS): HPFS is supported by the National Institutes of Health

(P01 CA055075, UM1 CA167552, U01 CA167552, R01 CA137178, R01 CA151993, R35

CA197735, K07 CA190673, and P50 CA127003), NHS by the National Institutes of Health (R01 CA137178, P01 CA087969, UM1 CA186107, R01 CA151993, R35 CA197735, K07

CA190673, and P50 CA127003) and PHS by the National Institutes of Health (R01 CA042182). Hawaii Adenoma Study: NCI grants R01 CA72520.

HCES-CRC: the Hwasun Cancer Epidemiology Study–Colon and Rectum Cancer (HCES-CRC; grants from Chonnam National University Hwasun Hospital, HCRI15011-1).

Kentucky: This work was supported by the following grant support: Clinical Investigator Award from Damon Runyon Cancer Research Foundation (CI-8); NCI R01CA136726.

LCCS: The Leeds Colorectal Cancer Study was funded by the Food Standards Agency and Cancer Research UK Programme Award (C588/A19167).

MCCS cohort recruitment was funded by VicHealth and Cancer Council Victoria. The MCCS was further supported by Australian NHMRC grants 509348, 209057, 251553 and 504711 and by infrastructure provided by Cancer Council Victoria. Cases and their vital status were ascertained through the Victorian Cancer Registry (VCR) and the Australian Institute of Health and Welfare (AIHW), including the National Death Index and the Australian Cancer Database.

MEC: National Institutes of Health (R37 CA54281, P01 CA033619, and R01 CA063464).

MECC: This work was supported by the National Institutes of Health, U.S. Department of Health and Human Services (R01 CA81488 to SBG and GR).

MSKCC: The work at Sloan Kettering in New York was supported by the Robert and Kate Niehaus Center for Inherited Cancer Genomics and the Romeo Milio Foundation. Moffitt: This work was supported by funding from the National Institutes of Health (grant numbers R01 CA189184, P30 CA076292), Florida Department of Health Bankhead-Coley Grant 09BN-13, and the University of South Florida Oehler Foundation. Moffitt contributions were supported in part by the Total Cancer Care Initiative, Collaborative Data Services Core, and Tissue Core at the H. Lee Moffitt Cancer Center & Research Institute, a National Cancer Institute-designated Comprehensive Cancer Center (grant number P30 CA076292).

NCCCS I & II: We acknowledge funding support for this project from the National Institutes of Health, R01 CA66635 and P30 DK034987.

NFCCR: This work was supported by an Interdisciplinary Health Research Team award from the Canadian Institutes of Health Research (CRT 43821); the National Institutes of Health, U.S. Department of Health and Human Serivces (U01 CA74783); and National Cancer Institute of Canada grants (18223 and 18226). The authors wish to acknowledge the contribution of Alexandre Belisle and the genotyping team of the McGill University and Génome Québec Innovation Centre, Montréal, Canada, for genotyping the Sequenom panel in the NFCCR samples. Funding was provided to Michael O. Woods by the Canadian Cancer Society Research

Institute.

NSHDS: Swedish Research Council; Swedish Cancer Society; Cutting-Edge Research Grant and other grants from the County Council of Västerbotten, Sweden; Wallenberg Centre for Molecular Medicine at Umeå University; Lion’s Cancer Research Foundation at Umeå University; the Cancer Research Foundation in Northern Sweden; and the Faculty of Medicine, Umeå University, Umeå, Sweden.

OFCCR: National Institutes of Health, through funding allocated to the Ontario Registry for Studies of Familial Colorectal Cancer (U01 CA074783); see CCFR section above. Additional funding toward genetic analyses of OFCCR includes the Ontario Research Fund, the Canadian Institutes of Health Research, and the Ontario Institute for Cancer Research, through generous support from the Ontario Ministry of Research and Innovation.

OSUMC: OCCPI funding was provided by Pelotonia and HNPCC funding was provided by the NCI (CA16058 and CA67941).

PLCO: Intramural Research Program of the Division of Cancer Epidemiology and Genetics and supported by contracts from the Division of Cancer Prevention, National Cancer Institute, NIH,

PMH: National Institutes of Health (R01 CA076366 to P.A. Newcomb).

SEARCH: The University of Cambridge has received salary support in respect of PDPP from the NHS in the East of England through the Clinical Academic Reserve. Cancer Research UK (C490/A16561); the UK National Institute for Health Research Biomedical Research Centres at the University of Cambridge.

SELECT: The Selenium and Vitamin E Cancer Prevention Trial (SELECT) was supported by the National Cancer Institute of the National Institutes of Health under Award Numbers UM1CA182883 and U10CA37429.

SMS: This work was supported by the National Cancer Institute (grant P01 CA074184 to J.D.P. and P.A.N., grants R01 CA097325, R03 CA153323, and K05 CA152715 to P.A.N., and the

National Center for Advancing Translational Sciences at the National Institutes of Health (grant KL2 TR000421 to A.N.B.-H.)

The Swedish Low-risk Colorectal Cancer Study: The study was supported by grants from the Swedish research council; K2015-55X-22674-01-4, K2008-55X-20157-03-3, K2006-72X-

20157-01-2 and the Stockholm County Council (ALF project).

Swedish Mammography Cohort and Cohort of Swedish Men: This work is supported by the Swedish Research Council /Infrastructure grant, the Swedish Cancer Foundation, and the Karolinska Institute´s Distinguished Professor Award to Alicja Wolk.

UK Biobank: This research has been conducted using the UK Biobank Resource under Application Number 8614

VITAL: National Institutes of Health (K05 CA154337).

WHI: The WHI program is funded by the National Heart, Lung, and Blood Institute, National Institutes of Health, U.S. Department of Health and Human Services through contracts HHSN268201100046C, HHSN268201100001C, HHSN268201100002C, HHSN268201100003C, HHSN268201100004C, and HHSN271201100004C.

Acknowledgements:

ASTERISK: We are very grateful to Dr. Bruno Buecher without whom this project would not have existed. We also thank all those who agreed to participate in this study, including the patients and the healthy control persons, as well as all the physicians, technicians and students.

CLUE II: We thank Judith Hoffman-Bolton, Senior Research Program Coordinator, for her contributions to the conduct of CLUE and its participation in this study. We also thank the participants in CLUE.

COLON: the authors would like to thank the COLON investigators at Wageningen University & Research and the involved clinicians in the participating hospitals.

CORSA: We kindly thank all those who contributed to the screening project Burgenland against CRC. Furthermore, we are grateful to Doris Mejri and Monika Hunjadi for laboratory assistance.

CPS-II: The authors thank the CPS-II participants and Study Management Group for their invaluable contributions to this research. The authors would also like to acknowledge the contribution to this study from central cancer registries supported through the Centers for Disease Control and Prevention National Program of Cancer Registries, and cancer registries supported by the National Cancer Institute Surveillance Epidemiology and End Results program.

Czech Republic CCS: We are thankful to all clinicians in major hospitals in the Czech Republic, without whom the study would not be practicable. We are also sincerely grateful to all patients participating in this study.

DACHS: We thank all participants and cooperating clinicians, and Ute Handte-Daub, Utz Benscheid, Muhabbet Celik and Ursula Eilber for excellent technical assistance.

EDRN: We acknowledge all the following contributors to the development of the resource: University of Pittsburgh School of Medicine, Department of Gastroenterology, Hepatology and Nutrition: Lynda Dzubinski; University of Pittsburgh School of Medicine, Department of Pathology: Michelle Bisceglia; and University of Pittsburgh School of Medicine, Department of Biomedical Informatics.

EPICOLON: We are sincerely grateful to all patients participating in this study who were recruited as part of the EPICOLON project. We acknowledge the Spanish National DNA Bank, Biobank of Hospital Clínic–IDIBAPS and Biobanco Vasco for the availability of the samples. The work was carried out (in part) at the Esther Koplowitz Centre, Barcelona.

Harvard cohorts (HPFS, NHS, PHS): The study protocol was approved by the institutional review boards of the Brigham and Women’s Hospital and Harvard T.H. Chan School of Public Health, and those of participating registries as required. We would like to thank the participants and staff of the HPFS, NHS and PHS for their valuable contributions as well as the following state cancer registries for their help: AL, AZ, AR, CA, CO, CT, DE, FL, GA, ID, IL, IN, IA, KY, LA, ME, MD, MA, MI, NE, NH, NJ, NY, NC, ND, OH, OK, OR, PA, RI, SC, TN, TX, VA, WA, WY. The authors assume full responsibility for analyses and interpretation of these data.

Kentucky: We would like to acknowledge the staff at the Kentucky Cancer Registry.

LCCS: We acknowledge the contributions of Jennifer Barrett, Robin Waxman, Gillian Smith and Emma Northwood in conducting this study.

NCCCS I & II: We would like to thank the study participants, and the NC Colorectal Cancer Study staff.

NSHDS: We thank all participants in the NSHDS cohorts, the staff at the Department of Biobank Research, Umeå University, the staff at Biobanken norr, Västerbotten County Council, and the scientists managing the Northern Sweden Diet Database.

PLCO: The authors thank the PLCO Cancer Screening Trial screening center investigators and the staff from Information Management Services Inc and Westat Inc. Most importantly, we thank the study participants for their contributions that made this study possible.

PMH: The authors would like to thank the study participants and staff of the Hormones and Colon Cancer study.

SEARCH: We thank the SEARCH team.

SELECT: We thank the research and clinical staff at the sites that participated on SELECT study, without whom the trial would not have been successful. We are also grateful to the 35,533 dedicated men who participated in SELECT.

WHI: The authors thank the WHI investigators and staff for their dedication, and the study participants for making the program possible. A full listing of WHI investigators can be found at: http://www.whi.org/researchers/Documents%20%20Write%20a%20Paper/WHI%20Investigator

%20Short%20List.pdf
